# GRID – Genomics of Rare Immune Disorders: a highly sensitive and specific diagnostic gene panel for patients with primary immunodeficiencies

**DOI:** 10.1101/431544

**Authors:** Ilenia Simeoni, Olga Shamardina, Sri VV Deevi, Moira Thomas, Karyn Megy, Emily Staples, Deborah Whitehorn, Daniel Duarte, Rutendo Mapeta, Willem H Ouwehand, Christopher J Penkett, Paula Rayner-Matthews, Hannah Stark, Jonathan C Stephens, Kathleen E Stirrups, Ernest Turro, NIHR BioResource, Adrian Thrasher, Taco W Kuijpers, Kenneth GC Smith, Sinisa Savic, Siobhan O Burns, James E Thaventhiran, Hana Lango Allen

## Abstract

Primary Immune disorders affect 15,000 new patients every year in Europe. Genetic tests are usually performed on a single or very limited number of genes leaving the majority of patients without a genetic diagnosis. We designed, optimised and validated a new clinical diagnostic platform called GRID, Genomics of Rare Immune Disorders, to screen in parallel 279 genes, including 2015 IUIS genes, known to be causative of Primary Immune disorders (PID). Validation to clinical standard using more than 58,000 variants in 176 PID patients shows an excellent sensitivity, specificity. The customised and automated bioinformatics pipeline prioritises and reports pertinent Single Nucleotide Variants (SNVs), INsertions and DELetions (INDELs) as well as Copy Number Variants (CNVs). An example of the clinical utility of the GRID panel, is represented by a patient initially diagnosed with X-linked agammaglobulinemia due to a missense variant in the *BTK* gene with severe inflammatory bowel disease. GRID results identified two additional compound heterozygous variants in *IL17RC*, potentially driving the altered phenotype.

## Introduction

Primary immunodeficiency disorders (PID) are a heterogeneous group of conditions that lead to infection susceptibility, malignancies and autoimmunity. Although disease-causing variants in over 300 genes have been identified, only about 10-20% of patients receive a molecular diagnosis. Identification of a monogenic cause can inform gene-specific treatment, even for patients with incompletely penetrant variants^1,2^. Targeted high-throughput sequencing (HTS) gene panels offer a cost-effective approach for simultaneous screening of many disease-causing genetic alleles^3^.

Here we describe the design and validation to clinical standard of GRID (<u>G</u>enomics of <u>R</u>are <u>I</u>mmune <u>D</u>isorders, www.gridgenomics.org.uk), a targeted HTS panel for screening all known genetic causes of PID. We show that this panel covers 99% of both the full IUIS 2015 gene list^4^ and the Human Gene Mutation Database (HGMD) variants associated with PID. In addition to having a considerably wider gene coverage compared to other diagnostic PID panels^5,6^, our platform provides good coverage of GC-rich regions previously reported as difficult to capture. We provide sensitivity and specificity estimates by direct comparison of over 58,000 variants in 176 PID patients with both whole genome sequencing (WGS) and GRID panel data. We describe a customised and automated bioinformatics analysis pipeline to prioritise and report pertinent Single Nucleotide Variants (SNVs), INsertions and DELetions (INDELs) as well as Copy Number Variants (CNVs). Finally, we demonstrate the clinical utility of an unbiased gene screen on 86 PID patients who had not been genetically tested previously.

### GRID panel design and coverage optimisation

We designed the GRID panel to capture all genes known to contain PID-causing variants, including the complete 2015 IUIS classification^4^, currently 279 genes (Table S1). Regions targeted in the panel design are defined in Figure 1A, with an illustration of targeted and captured regions over *TBX1* gene. For prioritisation of variants for diagnostic reporting we defined regions of interest (ROI) as: coding or lincRNA exons plus 15bp either side; and non-coding PID-associated HGMD DM variants. The validation of the panel described henceforth is based on the ROI.

**Figure 1.**
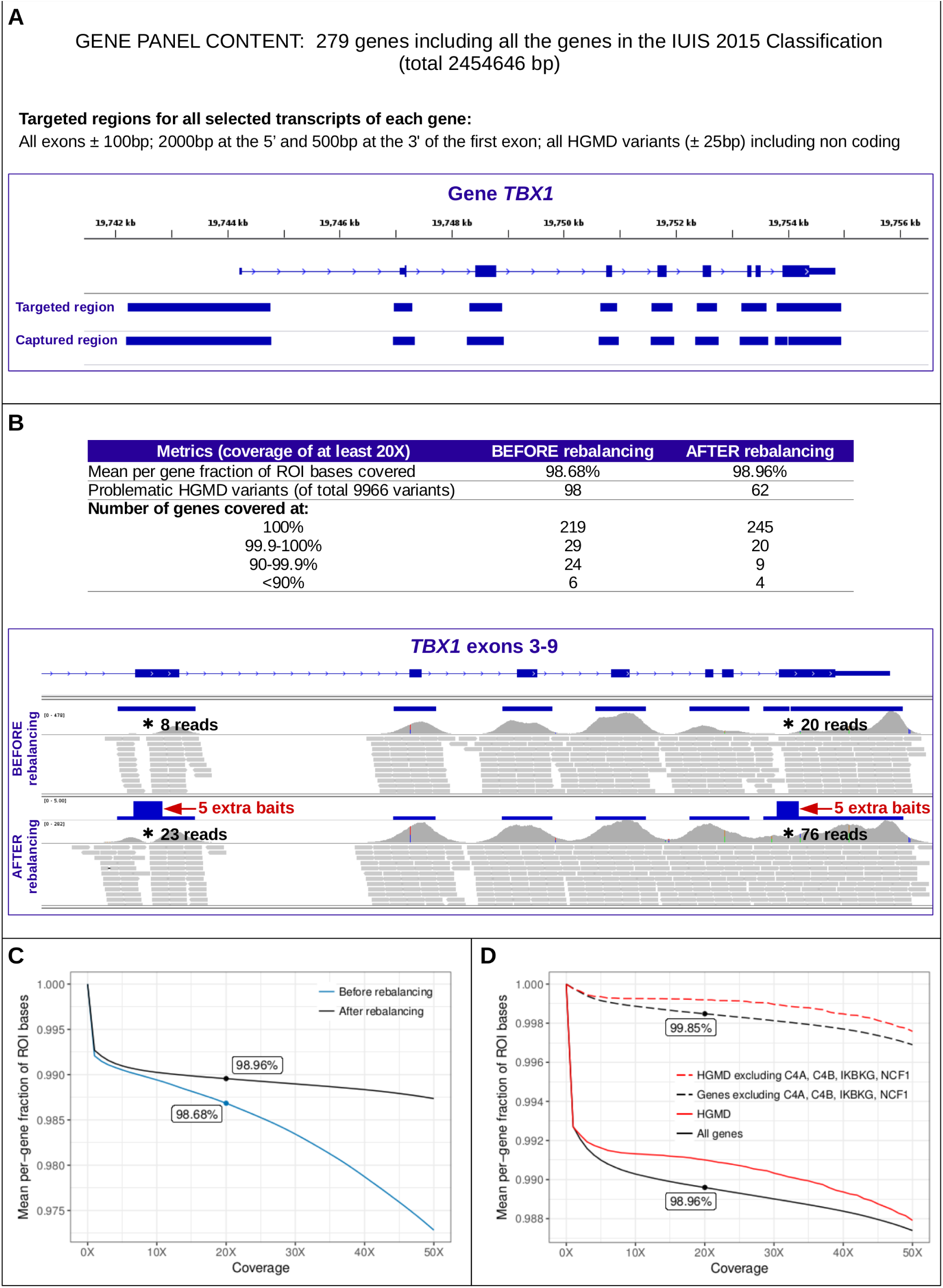
GRID panel design and coverage optimisation. (A) GRID panel content and example of targeted and captured regions in *TBX1* gene. (B) Improved coverage metrics and example of improved read depth in *TBX1*. (C) Mean per-gene fraction of ROI bases before and after rebalancing. (D) Mean per-gene fraction of ROI bases in all genes (solid black line) and HGMD variants (solid red line) in 176 samples; the same metrics excluding problematic genes (dashed lines). ROI = regions of interest.

To optimise panel coverage, we initially sequenced 59 samples and identified regions of sub-optimal (<20 reads) coverage in at least 5% of samples (problematic regions). We increased the number of probes in those regions through a process known as bait rebalancing (Figure 1B and Supplementary Methods). We then sequenced 176 samples from PID patients, and use these to show the coverage improvement by bait rebalancing and to report our final coverage metrics. We capture 643,088 of the 655,348 bases representing the ROI by at least 20 reads, corresponding to the average per gene fraction of ROI coverage of 98.96% (median 100%) (Figures 1B and 1C). The number of HGMD variants covered at ≥20X increased from 99.0% to 99.4% (Figure 1B). Overall, the mean coverage of the ROI across the 176 samples was 215X (min 148, max 314).

An example of improved coverage in one gene is shown in Figure 1B. Furthermore, in a number of genes previously reported to be difficult to capture, such as *JAK3*^*7*,8^, we obtained 100% coverage of at least 20X for all coding bases (Figure S1). Overall, all but 13 genes have >99.9% ROI coverage of ≥20X (Figure S2). Regions of systematically low coverage are primarily restricted to 4 genes (*C4A*, *C4B*, *IKBKG* and *NCF1*, see Figures 1D and S2), which have a high proportion of non-uniquely mapped reads owing to regions of high sequence homology with these genes elsewhere in the genome. This is a general limitation of short-read sequencing technology that cannot be overcome by bait rebalancing.

### GRID bioinformatics analysis and validation

We further developed the automated bioinformatics pipeline previously reported by Simeoni et al. ^9^ Improvements were made in: pertinent variant prioritisation within the ROI (Figure 2A); gene coverage quality control for each sample; and the CNV calling algorithm (Figure 2B and Supplementary Methods).

**Figure 2.**
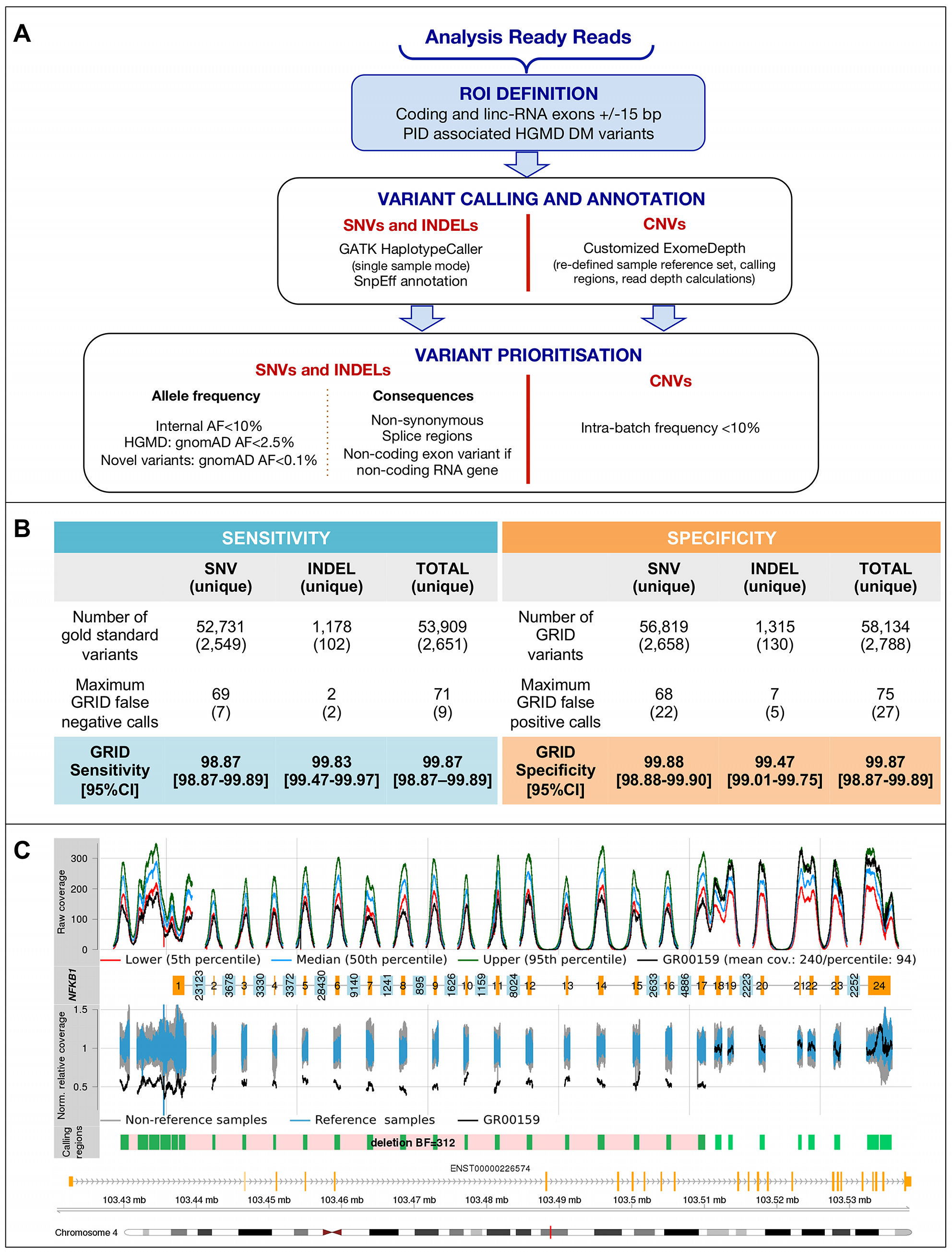
(A) GRID data processing, variant calling and prioritisation. (B) Sensitivity and Specificity of SNV and INDEL detection. (C) CNV plot example showing *NFKB1* heterozygous deletion call (pink block in the “Calling regions” track) made by the CNV pipeline automatically combining individual calls over exons 1-17 (green bars). “Norm. relative coverage” track visually confirms ~50% relative coverage in this sample (black lines) compared to reference samples (blue lines). Full explanation of the plot is provided in Supplementary Methods.

We assessed the sensitivity and specificity of the GRID panel using the 176 PID patient samples previously analysed by WGS, of which 53 patients had at least one pathogenic variant identified. We based these analyses on the ROI of the 279 PID genes. Before computing sensitivity and specificity metrics, we attempted to resolve all discrepant calls by Sanger sequencing to create a gold standard set of variant calls. Based on 52,731 SNV and 1,178 INDEL gold standard calls across all 176 samples, we obtained sensitivities of 99.87% (95% CI 98.87%−99.89%) and 99.83% (95% CI 99.47%−99.97%), respectively. The specificity values obtained for the 56,819 SNVs and 1,315 INDELs called by the GRID platform where the WGS data had ≥20X coverage and pass filter call over those positions (even if called as homozygous reference), were 99.88% (95% CI 98.88%−99.90%) and 99.47% (95% CI 99.01%−99.75%), respectively. All 69 pertinent variants reported across the 53 patients based on WGS data were correctly called and prioritised by the GRID pipeline, including 6 large deletions.

Intra-run reproducibility of the platform was assessed by sequencing three library preparations of the same sample on a single plate. The average pairwise genotype concordance of within ROI was 98.9%. Another set of three samples with three library preparations each was used to assess inter-run reproducibility, and the average pairwise ROI genotype concordance rates across three plates was 97.3% (Supplementary Methods and Table S2). Within the exonic and HGMD regions the intra- and inter-run concordances were even higher, at 99.2% and 98.9%, respectively. Together, these data show that the GRID panel generates highly reproducible variant calls, especially within the regions known to contain PID-causing variants.

We made further use of the WGS data to optimise the specificity of automated CNV calls, which has been rarely performed in previously described diagnostic gene panels. The GRID CNV pipeline identified 11 deletions and 21 duplications in 176 PID samples. WGS data confirmed 9 deletions and 10 duplications, ranging from 1 exon to multi-gene CNVs. All false calls had a lower confidence as measured by the Bayes factor statistic, allowing us to set a threshold above which to report CNVs (Figure S3). The 9 reliable deletion calls spanned 18 genes (single gene calls: *ARPC1B*, *CFHR4*, *DOCK8*, *IGKC*, *LRBA*, *NFKB1*, *TBX1* and one multi-gene call: *ATM*, *CD3D*, *CD3E*, *CD3G*, *CTSC*, *IL10RA*, *IL18*, *MRE11*, *SLC37A4*, *TIRAP*, *ZBTB16*), and 10 duplication calls spanned 9 genes (*CD8A*, *DOCK8*, *IKBKB*, *LYST*, *MASP2*, *MCM4*, *PRKDC*, *STAT2*, *UNG*). Each CNV can be visually inspected in an automatically generated plot (Figure 2B).

### Clinical application

We assessed the clinical utility of the GRID panel using new referrals of 86 PID patients without a genetic diagnosis. The automated prioritisation of variants generated an average of 5 variants (4.3 SNVs, 0.5 INDELs and 0.2 CNVs) per sample. All prioritised variants were reviewed in a Multi-Disciplinary Team meeting and categorised as clearly pathogenic, likely pathogenic or of uncertain significance, and for their contribution to the patient’s phenotype. This resulted in a genetic diagnosis for 10 of patients (11.6%). One patient with previously diagnosed Bruton’s agammaglobulinaemia and inflammatory bowel disease had disease-causing variants in both *IL17RC (p.Arg378* and p.Arg751Serfs*14)* and *BTK (p.Glu479Lys)*. Conventionally, this patient would have had a Sanger sequencing test of the *BTK* gene as a confirmation of the clinical diagnosis, albeit with an unusual presentation. The unbiased GRID approach identified an additional variant in a second gene that explains what was initially thought to be an expanded *BTK* phenotype.

In conclusion, we report the design, optimisation and validation of a new HTS panel for PID patients. We show that the GRID panel has excellent sensitivity and specificity for SNVs, INDELs and CNVs, and overcomes previously encountered challenges with coverage of GC-rich regions harbouring clinically relevant variants. We demonstrate its potential as a first-line, unbiased genetic test to provide a conclusive molecular diagnosis for patients suspected of having a highly penetrant genetic basis of their immune disorder.

## Supporting information

Online Repository

## Acknowledgements

Funding for the project was provided by the National Institute for Health Research (NIHR, grant number RG65966). J.E.D.T. is supported by the MRC (RG95376 and MR/L006197/1). A.J.T. is supported by the Wellcome Trust (104807/Z/14/Z) and the NIHR Biomedical Research Centre at Great Ormond Street Hospital for Children NHS Foundation Trust and University College London.

