## Supplementary material for "GRID – Genomics of Rare Immune Disorders: a highly sensitive and specific diagnostic gene panel for patients with primary immunodeficiencies": Online Repository

**Simeoni I *et al.***

1. Supplementary Methods
2. Supplementary References
3. Supplementary Tables
4. Supplementary Figures

### 1. Supplementary Methods

#### 1.1 Library preparation, enrichment and sequencing

Genomic DNA is extracted from blood, saliva or archived DNA as previously described[1]. The DNA quality is checked by gel electrophoresis and by two independent measurements of DNA concentration by Qubit (Life Technologies). DNA samples are processed in batches of 96 samples. 500ng of each DNA sample is fragmented using Covaris E220 (Covaris Inc., Woburn, MA, USA) to obtain an average size of 350 base pair DNA fragments. DNA samples are processed using the ROCHE KAPA HTP Library Preparation kit (Roche Diagnostics Ltd., Burgess Hill, UK). Six DNA libraries are captured using one reaction of ROCHE NimbleGen SeqCap GRID capture of 3Mb (ROCHE NimbleGen, Inc. Madison, WI USA). The capture step uses in-solution biotinylated DNA oligos (baits) to target selected regions of interest in the genome. Final libraries are quantified and 96 samples are pooled and sequenced in one lane of Illumina Hiseq 4000 sequencer, 150 base pair (bp) paired-end (PE) run.

#### 1.2 Problematic regions

A base was defined as problematic if more than 5% of samples had fewer than 20 reads with mapping quality greater than 20. Regions comprising consecutive problematic bases are shown in gene coverage plots that are generated for each batch of 96 samples processed. Figure S1 shows an example of a coverage plot in which problematic regions are visualised in red.

#### 1.3 Target capture optimisation

The optimisation of the capture design was performed implementing a 5x targeted replication. Probes targeting regions identified as problematic in the initial run received 5 probes for every probe in non-problematic regions. After the capture optimisation, 246 genes (including the *APOL1* gene that was added to the panel after the initial run) had at least 20X coverage of each base within their Regions Of Interest (ROI, defined as exons +15bp either side, plus non-coding HGMD DM variants). The details of exons and HGMD variants not covered at 20X, based on 176 samples, are shown Table S1. Genes with <100% ROI coverage at 20X are shown in Figure S2.

#### 1.4. SNV and INDEL calling and prioritisation

Single nucleotide variants (SNVs) and short insertions or deletions (INDELs) are called using GATK[5] 3.3 HaplotypeCaller in a single sample mode and filtered using the following VariantFiltration expressions "`MQ < 40.0 || QD < 2.0 || FS > 100.0`" for SNVs and "`FS > 200.0 || QD < 2.0 ||`

ReadPosRankSum < -20.0" for INDELs. Variants are then merged into a multi-sample VCF file. SNVs and INDELs are annotated with their predicted impact against Ensembl 75, presence in the human gene mutation database (HGMD[6]) version 2017.2, and their Minor Allele Frequency (MAF) in the Genome Aggregation Database (gnomAD[7]) release r.2.0.1 using SnpEff[8] 4.0.

Variants called within the ROI are then prioritised as follows: HGMD variants are retained if gnomAD AF is <2.5%; all other variants must have gnomAD AF <0.1%, have <4 alternate alleles (to guard against errors in repetitive regions), and be within one of the lincRNA genes (*RMRP*, *TERC*) or have a predicted moderate or high impact on translation of one of the GRID transcripts according to SnpEff. In addition, any variants with an internal AF >= 10% are removed as systematic artefacts.

### 1.5. CNV calling

CVNs are called using a custom pipeline based on the ExomeDepth[9] R-package (version 1.1.10). ExomeDepth makes a copy number gain or loss call by comparing the read depth in a specified genomic interval in a sample to that of an optimised reference set of other samples in the same batch. Our customisation specifically defines the reference set in order to eliminate false negative and positive calls that occur when the automatically chosen set includes too few samples for comparison. We also take into account sample relatedness within a batch: related samples and replicates are excluded from mutual reference sets and only an unrelated pool of samples is used for a reference set.

In order to detect smaller CNVs within large exons, we specify the target genomic intervals to be no longer than 500bp. Additionally, we modified the ExomeDepth read counting to avoid inflation caused by reads overlapping two adjacent regions. CNVs observed in more than 10% of samples within a batch are filtered out as technical artefacts or common CNVs.

ExomeDepth is most suitable for calling rare CNVs. A small number of regions lacking uniquely mapped reads due to gene homology are excluded from CNV calling so as not to bias the optimised reference sets and hence calling in the other regions. The excluded regions range from one exon to entire genes and are indicated in Table S1.

WGS data allowed us to assess the reliability of the GRID CNV calls. One of the 176 samples had an excessive number of one-exon deletions across different chromosomes, and was excluded from further analysis as a technical outlier. The rest of the samples had a total of 32 CNV calls, and we looked for evidence of these in the WGS data. The result of the comparison with WGS data is shown in Figure S3. All GRID calls that were supported by WGS data had Bayes factor (BF) higher than 20. This observation led us to set the BF>20 as a threshold for automatically reporting CNVs (although other less confident calls can nevertheless be inspected). In each case, the total length of the region over which the CNV call was made was greater than 1000 bases, suggesting that this may be the lower limit for the length of a CNV that can be reliably called.

#### 1.5a CNV plots

In order to visually assess CNV calls, we use Gviz[10] R package (version 1.22.3) to generate automated multi-track plots of raw and normalised coverage profiles of each sample for each gene where a CNV call over one or more exons was made (Figure 2B). The tracks of each CNV plot show the following: the lower, median and upper coverage percentiles of the exonic regions in the gene for all samples in the batch, and the raw coverage for the sample with the CNV call; numbered exons of the gene (yellow), and, if applicable, the number of intronic bases cut out from the plot in order to reduce its width (light blue); the normalised relative coverage for all samples in the batch (see below), where the sample with the CNV call is shown in black, its reference samples in blue, and other samples in the batch in grey; the custom-defined genomic intervals over which the calls were made (green) and the Bayes factor for the CNV call; and a to-scale representation of the Ensembl transcript used and genomic coordinates.

#### 1.5b Normalised coverage calculation

Relative coverage of a base in a sample was defined as the raw coverage of that base divided by the mean coverage of all bases in that sample. We normalised this relative coverage across all samples in a batch by dividing the relative base coverage in a sample by the mean of relative base coverages across all samples except the sample of interest. When assessing CNV calls in autosomal chromosomes, the expected normalised relative coverage is 1 for normal copy number; 0.5 for heterozygous deletion; and 1.5 for heterozygous duplication. In Figure 2B the sample with the partial *NFKB1* deletion has a drop in normalised coverage in exons 1-17 consistent with a heterozygous deletion.

#### 1.6 Intra-run and inter-run reproducibility

For intra-run assessment, three library preparations (L1, L2, L3) of a single sample (S1) were sequenced within the same 96-sample multiplex plate (P1). We then performed pairwise comparisons of genotype calls for the three L1-L2, L1-L3 and L2-L3 combinations and calculated the average intra-run concordance (Table S2). For inter-run assessment, three library preparations of another three samples (S2, S3, S4) were each sequenced in three time-independent runs of separate plates (P1, P2, P3). For each of the samples we then compared pairwise genotype calls across P1-P2, P1-P3 and P2-P3, and calculated the average per-sample inter-run concordance.

Calls were considered to be concordant if both the genotype and genomic location were exactly the same in each pairwise comparison. The concordance rates shown in Table S2 are for ROIs, which sometimes include homopolymeric intronic regions within the 15bp of the exons that can have INDEL calls with inconsistently assigned genomic locations. If these differ even by a single base, the calls will be called as discordant. When we restrict the comparisons to exons, splice sites and non-coding HGMD variants only, the intra-run and inter-run reproducibility increases from 98.9% to 99.2%, and from 97.3% to 98.9%, respectively.

### 3. Supplementary Tables

**Table S1. GRID panel genes and coverage.** For each gene the specific transcript used for the panel design is listed. The coverage metric refers to the percentage of bases covered by at least 20 reads within the ROI used for variant prioritisation.

**Table S2. Intra- and inter-run reproducibility of variant calls in (a) ROI and (b) exonic+splice site regions and HGMD sites.** Intra-run agreement was assessed by sequencing sample S1 three times on the same plate P1, using three library preparations L1, L2 and L3. Inter-run agreement was based on sequencing three library preparations (L1, L2, L3) of samples S2, S3 and S4 across plates P1, P2 and P3.

#### (a) ROI

| | | Comparison 1 | | Comparison 2 | | Comparison 3 | | Mean of Comparison 1,2,3 | |
|---|---|---|---|---|---|---|---|---|---|
| | Sample | comparison (Library/Plate) | concordance (%) | comparison (Library/Plate) | concordance (%) | comparison (Library/Plate) | concordance (%) | | |
| Intra-run (N variants) | S1 (406) | P1 L1-L2 | 98.77 | P1 L1-L3 | 98.52 | P1 L2-L3 | 99.26 | 98.85 | |
| Inter-run (N variants) | S2 (432) | L1.P1-L2.P2 | 97.69 | L1.P1-L3.P3 | 98.40 | L2.P2-L3.P3 | 95.37 | 97.15 | 97.29 |
| | S3 (419) | | 97.14 | | 97.85 | | 97.85 | 97.61 | |
| | S4 (417) | | 96.88 | | 97.84 | | 96.64 | 97.12 | |

#### (b) Exon + splice site + HGMD

| | | Comparison 1 | | Comparison 2 | | Comparison 3 | | Mean of Comparison 1,2,3 | |
|---|---|---|---|---|---|---|---|---|---|
| | Sample | comparison (Library/Plate) | concordance (%) | comparison (Library/Plate) | concordance (%) | comparison (Library/Plate) | concordance (%) | | |
| Intra-run (N variants) | S1 (303) | P1 L1-L2 | 99.34 | P1 L1-L3 | 99.01 | P1 L2-L3 | 99.34 | 99.23 | |
| Inter-run (N variants) | S2 (320) | L1.P1-L2.P2 | 99.37 | L1.P1-L3.P3 | 99.69 | L2.P2-L3.P3 | 99.06 | 99.37 | 98.94 |
| | S3 (308) | | 98.70 | | 99.03 | | 99.03 | 98.92 | |
| | S4 (319) | | 98.12 | | 98.75 | | 98.75 | 98.54 | |

### 4. Supplementary Figures

**Figure S1. Example of a coverage plot automatically generated for each gene for a batch of 96 samples processed.** Regions with less than 20 reads in more than 5% of samples are highlighted in red below the raw coverage track. In this case two problematic regions were identified in the 3'UTR of *JAK3*. Prob. regions: problematic regions.

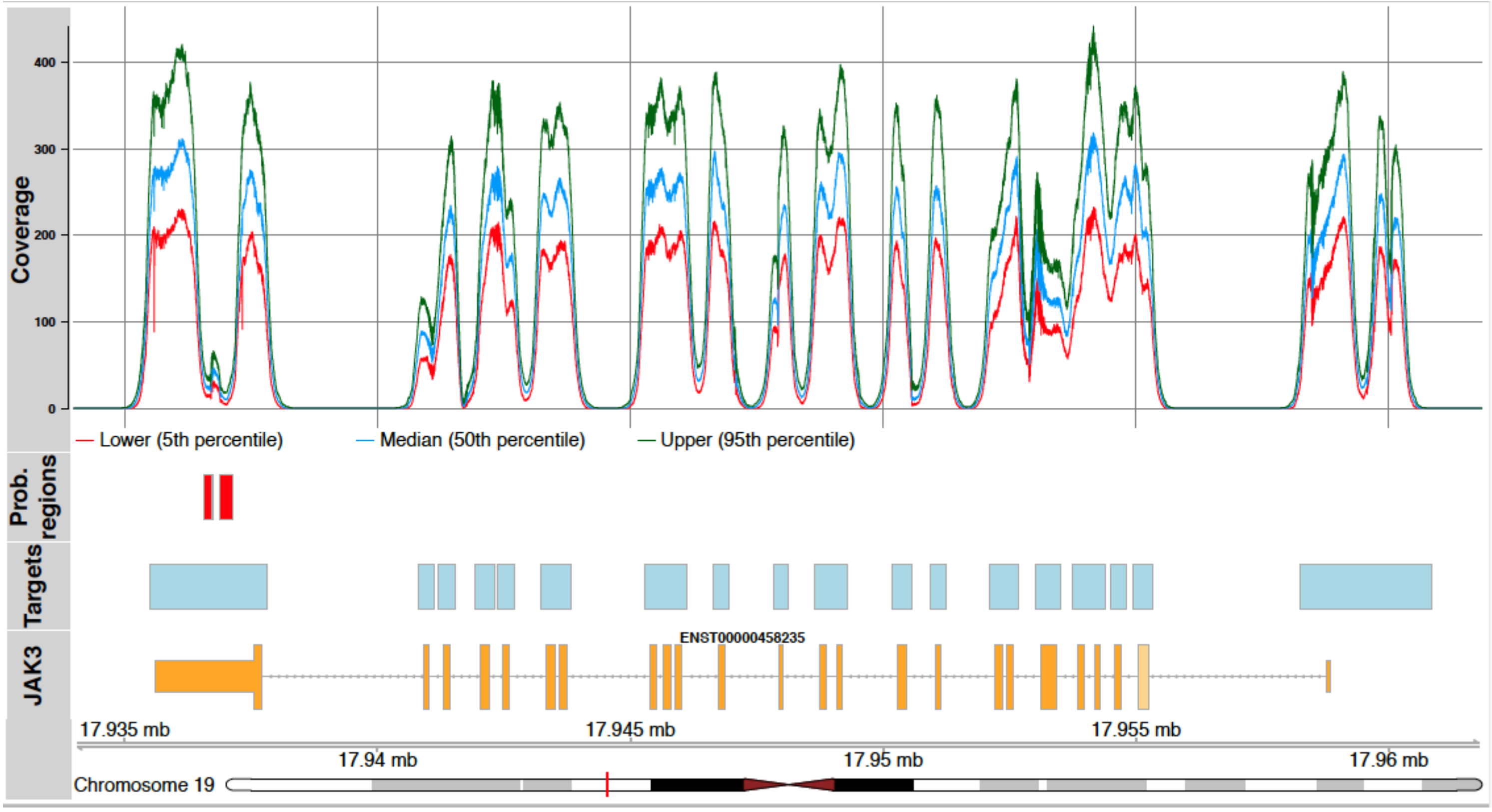

**Figure S2. GRID platform ROI coverage per gene for 33 genes with <100% coverage.** The remaining 246 genes with complete 100% ROI covered by at least 20X are excluded from this plot (see Table S1 for the full gene list and coverage details). The intervals shown in the plot exclude the left interval boundary.

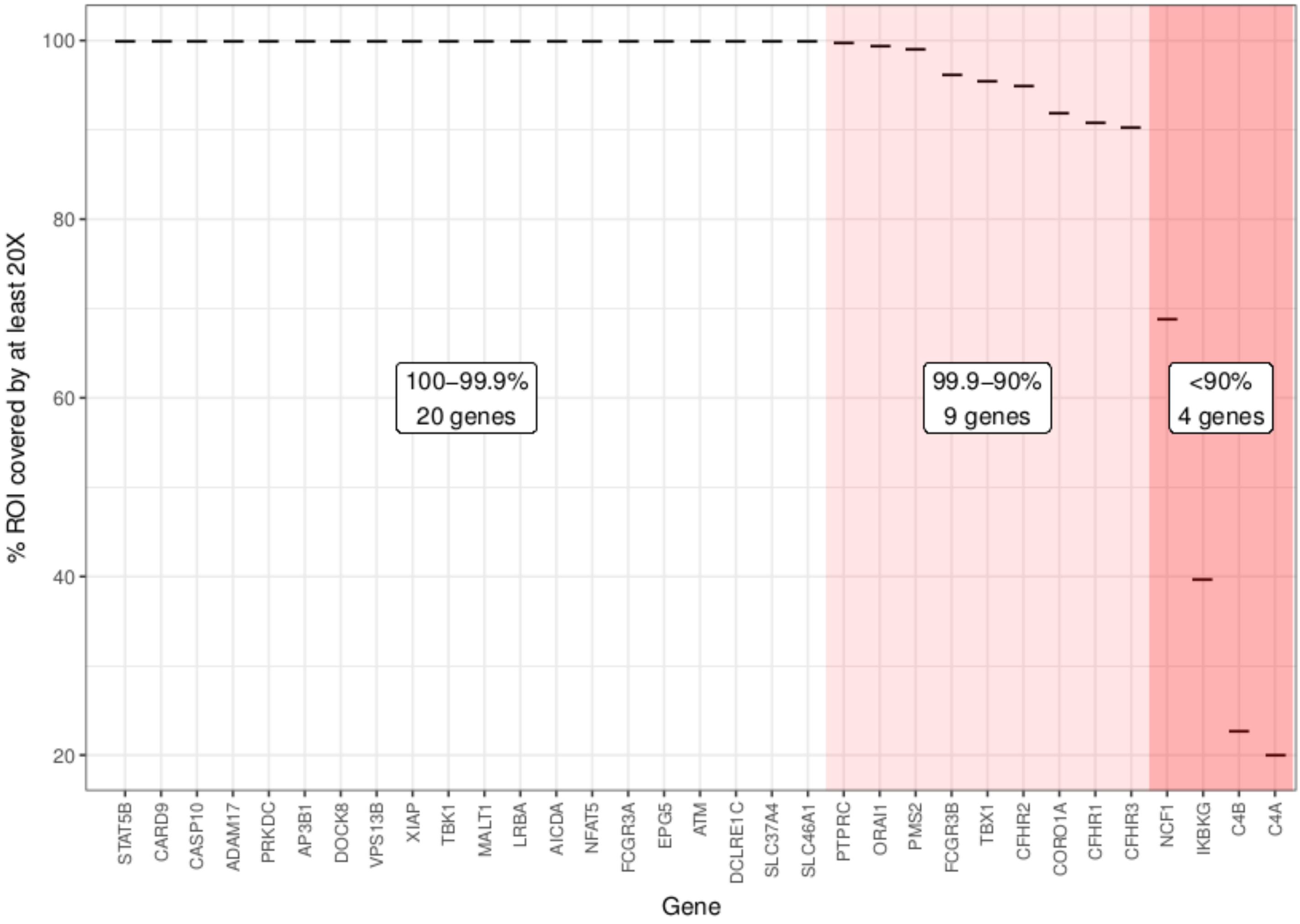

**Figure S3. Comparison of GRID CNV calls with WGS data.** The highlighted area of low Bayes factors and short CNVs corresponds to CNVs deemed false positive after manual inspection of WGS IGV plots for the region containing the putative GRID call. All confirmed GRID calls had Bayes factor >20 allowing us to set this as a threshold above which CNV calls are automatically reported. Note the logarithmic scale of both axes.

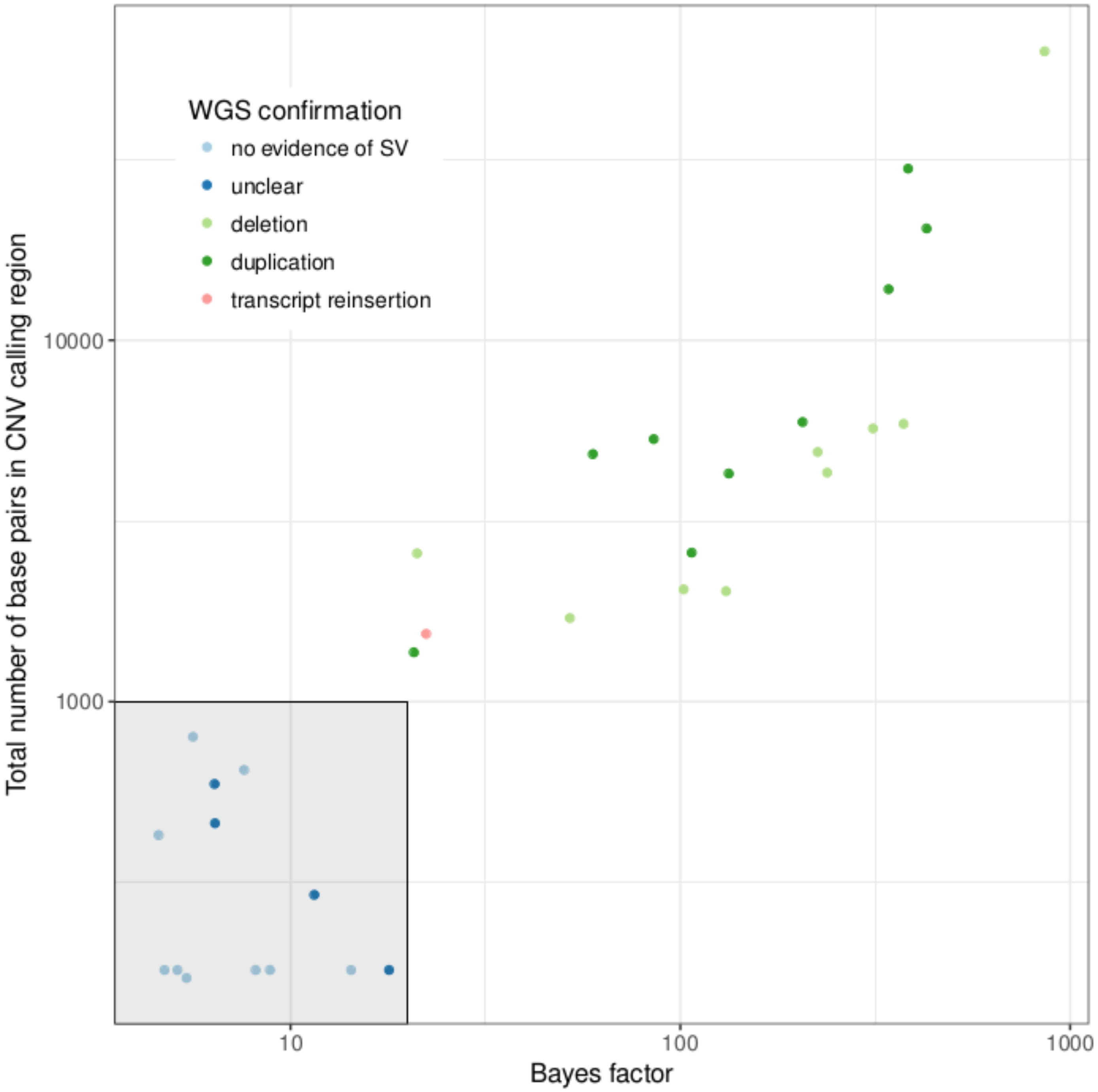
