## Supplementary Table S1 for "GRID – Genomics of Rare Immune Disorders: a highly sensitive and specific diagnostic gene panel for patients with primary immunodeficiencies"

| HGNC Gene Symbol | Transcript (RefSeq) | Transcript (Ensembl) | Locus Reference Genomic (LRG) ID | % ROI covered by at least 20X | Problematic exons | # HGMD variants in problematic ROI regions | Regions excluded from CNV calling |
|---|---|---|---|---|---|---|---|
| ACD | NM_001082486.1 | ENST00000219251 | | 100 | | | |
| ACP5 | NM_001111034.2 | ENST00000592828 | | 100 | | | |
| ACTB | NM_001101.3 | ENST00000331789 | LRG_132 | 100 | | | |
| ADA | NM_000022.2 | ENST00000372874 | LRG_16 | 100 | | | |
| ADA2 | NM_001282227.1 | ENST00000399839 | | 100 | | | |
| ADAM17 | NM_003183.4 | ENST00000310823 | | 99.9996 | | | |
| ADAR | NM_001111.4 | ENST00000368474 | | 100 | | | |
| AICDA | NM_020661.2 | ENST00000229335 | LRG_17 | 99.997 | | | |
| AIRE | NM_000383.3 | ENST00000291582 | LRG_18 | 100 | | | |
| AK2 | NM_013411.3 | ENST00000354858 | LRG_133 | 100 | | | |
| AP3B1 | NM_001271769 | ENST00000519295 | LRG_170 | 99.9996 | | | |
| APOL1 | NM_145343.2 | ENST00000319136 | LRG_169 | 100 | | | |
| ARPC1B | NM_005720 | ENST00000252725 | | 100 | | | |
| ATM | NM_000051.3 | ENST00000278616 | LRG_135 | 99.9907 | | 1 (CS135658) | |
| B2M | NM_004048.2 | ENST00000558401 | | 100 | | | |
| BCL10 | NM_003921.4 | ENST00000370580 | | 100 | | | |
| BLM | NM_000057.2 | ENST00000355112 | LRG_20 | 100 | | | |
| BLNK | NM_013314.3 | ENST00000224337 | LRG_21 | 100 | | | |
| BLOC1S6 | NM_012388.3 | ENST00000220531 | LRG_883 | 100 | | | |
| BTK | NM_000061.2 | ENST00000308731 | LRG_128 | 100 | | | |
| C1QA | NM_015991.2 | ENST00000374642 | LRG_22 | 100 | | | |
| C1QB | NM_000491.3 | ENST00000314933 | LRG_23 | 100 | | | |
| C1QC | NM_001114101.1 | ENST00000374639 | LRG_24 | 100 | | | |
| C1R | NM_001733 | ENST00000542285 | | 100 | | | |
| C1S | NM_201442/NM_001734 | ENST00000328916 | LRG_25 | 100 | | | |
| C2 | NM_000063.4 | ENST00000299367 | LRG_26 | 100 | | | |
| C3 | NM_000064.2 | ENST00000245907 | LRG_27 | 100 | | | |
| C4A | NM_007293.2 | ENST00000428956 | LRG_137 | 20.0389 | 1-20, 22-24, 29-30, 32-41 | 5 (CD045879, CM094583, CD983970, CI931069, CI094587) | whole gene |
| C4B | NM_001002029.3 | ENST00000435363 | LRG_138 | 22.6978 | 1-19, 21-24, 31-41 | 4 (CM920135, CM940213, CD024038, CI094586) | whole gene |
| C5 | NM_001735.2 | ENST00000223642 | LRG_28 | 100 | | | |
| C6 | NM_000065.2 | ENST00000263413 | LRG_29 | 100 | | | |
| C7 | NM_000587.2 | ENST00000313164 | LRG_30 | 100 | | | |
| C8A | NM_000562.2 | ENST00000361249 | LRG_139 | 100 | | | |
| C8B | NM_000066.2 | ENST00000371237 | LRG_31 | 100 | | | |
| C8G | NM_000606.2 | ENST00000224181 | | 100 | | | |
| C9 | NM_001737.3 | ENST00000263408 | LRG_32 | 100 | | | |
| CARD11 | NM_032415.5 | ENST00000396946 | LRG_729 | 100 | | | |
| CARD14 | NM_024110.4 | ENST00000344227 | | 100 | | | |
| CARD9 | NM_052813.4 | ENST00000371732 | LRG_178 | 99.9997 | | | |
| CASP10 | NM_032977.3 | ENST00000286186 | LRG_33 | 99.9997 | | | |
| CASP8 | NM_001228.4 | ENST00000358485 | LRG_34 | 100 | | | |
| CCBE1 | NM_133459.3 | ENST00000439986 | | 100 | | | |
| CD19 | NM_001770.5 | ENST00000538922 | LRG_35 | 100 | | | |
| CD247 | NM_198053.2 | ENST00000362089 | LRG_36 | 100 | | | |
| CD27 | NM_001242.4 | ENST00000266557 | LRG_357 | 100 | | | |
| CD3D | NM_000732.4 | ENST00000300692 | LRG_37 | 100 | | | |
| CD3E | NM_000733.3 | ENST00000361763 | LRG_38 | 100 | | | |
| CD3G | NM_000073.2 | ENST00000532917 | LRG_39 | 100 | | | |
| CD40 | NM_001250.4 | ENST00000372285 | LRG_40 | 100 | | | |
| CD40LG | NM_000074.2 | ENST00000370629 | LRG_141 | 100 | | | |
| CD46 | NM_002389.4 | ENST00000322875 | LRG_155 | 100 | | | |
| CD59 | NM_203330.2 | ENST00000395850 | LRG_41 | 100 | | | |
| CD79A | NM_001783.3 | ENST00000221972 | LRG_42 | 100 | | | |
| CD79B | NM_001039933.1 | ENST00000392795 | LRG_43 | 100 | | | |
| CD81 | NM_004356.3 | ENST00000263645 | LRG_142 | 100 | | | |
| CD8A | NM_001768.6 | ENST00000409511 | LRG_44 | 100 | | | |
| CEBPE | NM_001805.2 | ENST00000206513 | LRG_45 | 100 | | | |
| CFB | NM_001710.5 | ENST00000425368 | LRG_136 | 100 | | | |
| CFD | NM_001928.2 | ENST00000327726 | LRG_46 | 100 | | | |
| CFH | NM_000186.3 | ENST00000367429 | LRG_47 | 100 | | | exon 20 |
| CFHR1 | NM_002113.2 | ENST00000320493 | LRG_149 | 90.8573 | 1-6 | 1 (CM098960) | whole gene |
| CFHR2 | NM_005666.3 | ENST00000367415 | | 95.0427 | 1 | 0 | exon 1 |
| CFHR3 | NM_021023.5 | ENST00000367425 | LRG_175 | 90.4165 | 1-6 | 1 (CD098961) | whole gene |
| CFHR4 | NM_001201551.1 | ENST00000367416 | | 100 | | | exon 9 |
| CFHR5 | NM_030787.3 | ENST00000256785 | LRG_227 | 100 | | | |
| CFI | NM_000204.3 | ENST00000394634 | LRG_48 | 100 | | | |
| CFP | NM_002621.2 | ENST00000247153 | LRG_129 | 100 | | | |

| | | | | | | | |
|---|---|---|---|---|---|---|---|
| CHD7 | NM_017780.2 | ENST00000423902 | LRG_176 | 100 | | | |
| CIITA | NM_000246.3 | ENST00000324288 | LRG_49 | 100 | | | |
| CLPB | NM_030813.5 | ENST00000294053 | | 100 | | | |
| COPA | NM_001098398.1 | ENST00000368069 | | 100 | | | |
| CORO1A | NM_007074.3 | ENST00000219150 | LRG_195 | 92.0077 | 11 | 0 | |
| CR2 | NM_001006658.2 | ENST00000367057 | LRG_348 | 100 | | | |
| CSF2RA | NM_001161529.1 | ENST00000432318 | LRG_186 | 100 | | | |
| CSF3R | NM_156039.3 | ENST00000373103 | LRG_144 | 100 | | | |
| CTLA4 | NM_005214.4 | ENST00000302823 | | 100 | | | |
| CTPS1 | NM_001905.3 | ENST00000372621 | | 100 | | | |
| CTSC | NM_001814.4 | ENST00000227266 | LRG_50 | 100 | | | |
| CXCR4 | NM_003467.2 | ENST00000409817 | LRG_51 | 100 | | | |
| CYBA | NM_000101.3 | ENST00000261623 | LRG_52 | 100 | | | |
| CYBB | NM_000397.3 | ENST00000378588 | LRG_53 | 100 | | | |
| DCLRE1B | NM_001319946.1 | ENST00000369563 | | 100 | | | |
| DCLRE1C | NM_001033855.2 | ENST00000378278 | LRG_54 | 99.96 | | 1 (CS119472) | |
| DKC1 | NM_001363.3 | ENST00000369550 | LRG_55 | 100 | | | |
| DNMT3B | NM_006892.3 | ENST00000328111 | LRG_56 | 100 | | | |
| DOCK2 | NM_004946.2 | ENST00000256935 | | 100 | | | |
| DOCK8 | NM_203447.3 | ENST00000453981 | LRG_196 | 99.9993 | | | |
| ELANE | NM_001972.2 | ENST00000590230 | LRG_57 | 100 | | | |
| EPG5 | NM_020964.2 | ENST00000282041 | | 99.9923 | | | |
| F12 | NM_000505.3 | ENST00000253496 | LRG_145 | 100 | | | |
| FADD | NM_003824.3 | ENST00000301838 | LRG_228 | 100 | | | |
| FAS | NM_000043.4 | ENST00000355740 | LRG_134 | 100 | | | |
| FASLG | NM_000639.1 | ENST00000367721 | LRG_58 | 100 | | | |
| FCGR3A | NM_001127593.1 | ENST00000367969 | LRG_60 | 99.9961 | | | whole gene |
| FCGR3B | NM_000570.4 | ENST00000531221 | | 96.3311 | 3 | 0 | whole gene |
| FCN3 | NM_003665.2 | ENST00000270879 | LRG_171 | 100 | | | |
| FERMT3 | NM_178443.2 | ENST00000279227 | LRG_180 | 100 | | | |
| FOXN1 | NM_003593.2 | ENST00000226247 | LRG_61 | 100 | | | |
| FOXP3 | NM_014009.3 | ENST00000376207 | LRG_62 | 100 | | | |
| FPR1 | NM_001193306 | ENST00000595042 | LRG_146 | 100 | | | |
| G6PC3 | NM_138387.3 | ENST00000269097 | LRG_182 | 100 | | | |
| GATA2 | NM_001145661.1 | ENST00000341105 | LRG_295 | 100 | | | |
| GFI1 | NM_005263.3 | ENST00000370332 | LRG_63 | 100 | | | |
| HAX1 | NM_006118.3 | ENST00000328703 | LRG_64 | 100 | | | |
| ICOS | NM_012092.3 | ENST00000316386 | LRG_65 | 100 | | | |
| IFIH1 | NM_022168.3 | ENST00000263642 | | 100 | | | |
| IFNGR1 | NM_000416.2 | ENST00000367739 | LRG_66 | 100 | | | |
| IFNGR2 | NM_005534.3 | ENST00000290219 | LRG_67 | 100 | | | |
| IGHM | | ENST00000390559 | | 100 | | | |
| IGKC | | ENST00000390237 | | 100 | | | |
| IGLL1 | NM_020070.2 | ENST00000330377 | LRG_69 | 100 | | | |
| IKBKB | NM_001556.2 | ENST00000520810 | | 100 | | | |
| IKBKG | NM_001099857.2 | ENST00000369609 | LRG_70 | 39.6866 | 3-10 | 41 (upon request) | exons 3-10 |
| IKZF1 | NM_006060.5 | ENST00000439701 | | 100 | | | |
| IL10 | NM_000572.2 | ENST00000423557 | | 100 | | | |
| IL10RA | NM_001558.3 | ENST00000227752 | LRG_151 | 100 | | | |
| IL10RB | NM_000628.4 | ENST00000290200 | LRG_152 | 100 | | | |
| IL12B | NM_002187.2 | ENST00000231228 | LRG_71 | 100 | | | |
| IL12RB1 | NM_005535.1 | ENST00000600835 | LRG_72 | 100 | | | |
| IL17A | NM_002190.2 | ENST00000340057 | | 100 | | | |
| IL17F | NM_052872.3 | ENST00000336123 | LRG_356 | 100 | | | |
| IL17RA | NM_014339.6 | ENST00000319363 | LRG_355 | 100 | | | |
| IL17RC | NM_153461.3 | ENST00000295981 | | 100 | | | |
| IL1RN | NM_173841.2 | ENST00000259206 | LRG_188 | 100 | | | |
| IL21 | NM_021803.3 | ENST00000264497 | | 100 | | | |
| IL21R | NM_181079.4 | ENST00000337929 | LRG_731 | 100 | | | |
| IL22 | NM_020525 | ENST00000538666 | | 100 | | | |
| IL2RA | NM_000417.2 | ENST00000379959 | LRG_73 | 100 | | | |
| IL2RG | NM_000206.2 | ENST00000374202 | LRG_150 | 100 | | | |
| IL36RN | NM_173170.1 | ENST00000393200 | LRG_730 | 100 | | | |
| IL7R | NM_002185.3 | ENST00000303115 | LRG_74 | 100 | | | |
| INO80 | NM_017553.2 | ENST00000361937 | | 100 | | | |
| IRAK4 | NM_016123.3 | ENST00000448290 | LRG_75 | 100 | | | |
| IRF7 | NM_001572.3 | ENST00000397566 | | 100 | | | |
| IRF8 | NM_002163.2 | ENST00000268638 | LRG_294 | 100 | | | |
| ISG15 | NM_005101.3 | ENST00000379389 | | 100 | | | |
| ITCH | NM_031483.4 | ENST00000262650 | LRG_354 | 100 | | | |
| ITGAM | NM_001145808.1 | ENST00000544665 | | 100 | | | |
| ITGB2 | NM_000211.3 | ENST00000397850 | LRG_76 | 100 | | | |
| ITK | NM_005546.3 | ENST00000422843 | LRG_189 | 100 | | | |
| JAGN1 | NM_032492.3 | ENST00000307768 | | 100 | | | |
| JAK3 | NM_000215.3 | ENST00000458235 | LRG_77 | 100 | | | |
| KRAS | NM_033360.3 | ENST00000256078 | LRG_344 | 100 | | | |
| LAMTOR2 | NM_014017.3 | ENST00000368305 | LRG_81 | 100 | | | |
| LCK | NM_005356.4 | ENST00000336890 | LRG_153 | 100 | | | |

| | | | | | | | |
|---|---|---|---|---|---|---|---|
| LIG1 | NM_000234.1 | ENST00000263274 | LRG_78 | 100 | | | |
| LIG4 | NM_002312.3 | ENST00000356922 | LRG_79 | 100 | | | |
| LPIN2 | NM_014646.2 | ENST00000261596 | LRG_174 | 100 | | | |
| LRBA | NM_006726.4 | ENST00000357115 | | 99.9972 | | | |
| LYST | NM_000081.3 | ENST00000389794 | LRG_143 | 100 | | | |
| MAGT1 | NM_032121.5 | ENST00000358075 | LRG_353 | 100 | | | |
| MALT1 | NM_006785.3 | ENST00000348428 | | 99.9983 | | | |
| MAP3K14 | NM_003954.4 | ENST00000344686 | | 100 | | | |
| MASP2 | NM_006610.3 | ENST00000400897 | LRG_82 | 100 | | | |
| MBL2 | NM_000242.2 | ENST00000373968 | LRG_154 | 100 | | | |
| MCM4 | NM_005914.3 | ENST00000262105 | | 100 | | | |
| MEFV | NM_000243.2 | ENST00000219596 | LRG_190 | 100 | | | |
| MOGS | NM_006302.2 | ENST00000233616 | | 100 | | | |
| MPO | NM_000250.1 | ENST00000225275 | LRG_84 | 100 | | | |
| MRE11 | NM_005591 | ENST00000323929 | LRG_85 | 100 | | | |
| MS4A1 | NM_152866.2 | ENST00000534668 | LRG_140 | 100 | | | |
| MSH6 | NM_000179.2 | ENST00000234420 | LRG_219 | 100 | | | |
| MTHFD1 | NM_005956.3 | ENST00000216605 | | 100 | | | |
| MVK | NM_000431.2 | ENST00000228510 | LRG_156 | 100 | | | |
| MYD88 | NM_001172567.1 | ENST00000417037 | LRG_157 | 100 | | | |
| NBN | NM_002485.4 | ENST00000265433 | LRG_158 | 100 | | | |
| NCF1 | NM_000265.5 | ENST00000289473 | LRG_87 | 68.9749 | 1, 5, 8-9, 11 | 6 (CS014965, CS014966, CM961019, CD101818, CD014969, CD094221) | exons 1, 3-11 |
| NCF2 | NM_000433.3 | ENST00000367535 | LRG_88 | 100 | | | |
| NCF4 | NM_013416.3 | ENST00000397147 | LRG_159 | 100 | | | |
| NFAT5 | NM_138714.3 | ENST00000432919 | | 99.9962 | | | |
| NFKB1 | NM_003998.3 | ENST00000226574 | | 100 | | | |
| NFKB2 | NM_001288724.1 | ENST00000369966 | | 100 | | | |
| NFKBIA | NM_020529.2 | ENST00000216797 | LRG_89 | 100 | | | |
| NHEJ1 | NM_024782.2 | ENST00000356853 | LRG_90 | 100 | | | |
| NHP2 | NM_017838.3 | ENST00000274606 | LRG_346 | 100 | | | |
| NLRC4 | NM_021209.4 | ENST00000404025 | | 100 | | | |
| NLRP12 | NM_144687.3 | ENST00000324134 | LRG_181 | 100 | | | |
| NLRP3 | NM_004895.4 | ENST00000336119 | LRG_197 | 100 | | | |
| NOD2 | NM_022162.1 | ENST00000300589 | LRG_177 | 100 | | | |
| NOP10 | NM_018648.3 | ENST00000328848 | LRG_345 | 100 | | | |
| NRAS | NM_002524.4 | ENST00000369535 | LRG_92 | 100 | | | |
| ORAI1 | NM_032790.3 | ENST00000330079 | LRG_93 | 99.4254 | 1 | 0 | |
| PARN | NM_002582.3 | ENST00000437198 | | 100 | | | |
| PGM3 | NM_001199917.1 | ENST00000506587 | | 100 | | | |
| PIK3CD | NM_005026.3 | ENST00000377346 | LRG_191 | 100 | | | |
| PIK3R1 | NM_181523.2 | ENST00000521381 | LRG_453 | 100 | | | |
| PLCG2 | NM_002661.4 | ENST00000359376 | LRG_376 | 100 | | | |
| PMS2 | NM_000535.5 | ENST00000265849 | LRG_161 | 99.1136 | 15 | 2 (CM1515443, CI146161) | exons 13-15 |
| PNP | NM_000270.3 | ENST00000361505 | LRG_91 | 100 | | | |
| POLE | NM_006231.3 | ENST00000320574 | LRG_789 | 100 | | | |
| PRF1 | NM_001083116.1 | ENST00000441259 | LRG_94 | 100 | | | |
| PRKCD | NM_006254.3 | ENST00000394729 | | 100 | | | |
| PRKDC | NM_006904.6 | ENST00000314191 | LRG_162 | 99.9996 | | | |
| PSMB8 | NM_004159.4 | ENST00000374882 | | 100 | | | |
| PSTPIP1 | NM_003978.3 | ENST00000558012 | LRG_172 | 100 | | | |
| PTPRC | NM_002838.4 | ENST00000442510 | LRG_95 | 99.8531 | 27 | 0 | |
| RAB27A | NM_004580.4 | ENST00000396307 | LRG_96 | 100 | | | |
| RAC2 | NM_002872.3 | ENST00000249071 | LRG_97 | 100 | | | |
| RAG1 | NM_000448.2 | ENST00000299440 | LRG_98 | 100 | | | |
| RAG2 | NM_000536.3 | ENST00000311485 | LRG_99 | 100 | | | |
| RBCK1 | NM_031229.2 | ENST00000356286 | LRG_728 | 100 | | | |
| RFX5 | NM_000449.3 | ENST00000290524 | LRG_101 | 100 | | | |
| RFXANK | NM_003721.3 | ENST00000303088 | LRG_102 | 100 | | | |
| RFXAP | NM_000538.3 | ENST00000255476 | LRG_103 | 100 | | | |
| RHOH | NM_004310.4 | ENST00000381799 | LRG_736 | 100 | | | |
| RMRP | NR_003051.3 | ENST00000602361 | LRG_163 | 100 | | | |
| RNASEH2A | NM_006397.2 | ENST00000221486 | LRG_278 | 100 | | | |
| RNASEH2B | NM_001142279.2 | ENST00000336617 | LRG_279 | 100 | | | |
| RNASEH2C | NM_032193.3 | ENST00000308418 | LRG_280 | 100 | | | |
| RNF168 | NM_152617.3 | ENST00000318037 | LRG_185 | 100 | | | |
| RNF31 | NM_017999.4 | ENST00000324103 | | 100 | | | |
| RORC | NM_005060.3 | ENST00000318247 | | 100 | | | |
| RPSA | NM_002295.4 | ENST00000301821 | LRG_735 | 100 | | | |
| RTEL1 | NM_016434.3 | ENST00000508582 | | 100 | | | |
| SAMHD1 | NM_015474.3 | ENST00000262878 | LRG_281 | 100 | | | |
| SART3 | NM_014706.3 | ENST00000228284 | | 100 | | | |
| SBDS | NM_016038.2 | ENST00000246868 | LRG_104 | 100 | | | |
| SEMA3E | NM_012431.2 | ENST00000307792 | | 100 | | | |

| | | | | | | |
|---|---|---|---|---|---|---|
| SERPING1 | NM 000062.2 | ENST00000278407 | LRG 105 | 100 | | |
| SH2D1A | NM 002351.4 | ENST00000371139 | LRG 106 | 100 | | |
| SH3BP2 | NM 001122681.1 | ENST00000503393 | | 100 | | |
| SLC29A3 | NM 018344.5 | ENST00000373189 | | 100 | | |
| SLC35C1 | NM 018389.4 | ENST00000314134 | LRG 107 | 100 | | |
| SLC37A4 | NM 001164277.1 | ENST00000357590 | LRG 187 | 99.9389 | 6 | 0 |
| SLC46A1 | NM 080669.3 | ENST00000440501 | LRG 183 | 99.9347 | 4 | 0 |
| SMARCAL1 | NM 014140.3 | ENST00000357276 | LRG 108 | 100 | | |
| SP110 | NM 080424.2 | ENST00000258381 | LRG 109 | 100 | | |
| SPINK5 | NM 006846.3 | ENST00000359874 | LRG 110 | 100 | | |
| STAT1 | NM 007315.3 | ENST00000361099 | LRG 111 | 100 | | |
| STAT2 | NM 005419.3 | ENST00000314128 | | 100 | | |
| STAT3 | NM 139276.2 | ENST00000264657 | LRG 112 | 100 | | |
| STAT5B | NM 012448.3 | ENST00000293328 | LRG 192 | 99.9998 | | |
| STIM1 | NM 003156.3 | ENST00000300737 | LRG 164 | 100 | | |
| STK4 | NM 006282.2 | ENST00000372806 | LRG 535 | 100 | | |
| STX11 | NM 003764.3 | ENST00000367568 | LRG 113 | 100 | | |
| STXBP2 | NM 006949.2 | ENST00000221283 | LRG 165 | 100 | | |
| TAP1 | NM 000593.5 | ENST00000354258 | LRG 166 | 100 | | |
| TAP2 | NM 000544.3 | ENST00000374899 | LRG 167 | 100 | | |
| TAPBP | NM 003190.4 | ENST00000426633 | LRG 114 | 100 | | |
| TAZ | NM 000116.3 | ENST00000299328 | LRG 131 | 100 | | |
| TBK1 | NM 013254.3 | ENST00000331710 | | 99.9986 | | |
| TBX1 | NM 080647.1 | ENST00000332710 | LRG 226 | 95.4855 | 3 | 0 |
| TCF3 | NM 003200.3 | ENST00000262965 | | 100 | | |
| TCN2 | NM 000355.2 | ENST00000215838 | LRG 116 | 100 | | |
| TERC | NR 001566.1 | ENST00000602385 | LRG 347 | 100 | | |
| TERT | NM 198253.2 | ENST00000310581 | LRG 343 | 100 | | |
| THBD | NM 000361.2 | ENST00000377103 | LRG 168 | 100 | | |
| TICAM1 | NM 182919.3 | ENST00000248244 | LRG 358 | 100 | | |
| TINF2 | NM 012461.2 | ENST00000267415 | | 100 | | |
| TLR3 | NM 003265.2 | ENST00000296795 | LRG 117 | 100 | | |
| TMC6 | NM 007267.6 | ENST00000322914 | LRG 118 | 100 | | |
| TMC8 | NM 152468.4 | ENST00000318430 | LRG 119 | 100 | | |
| TMEM173 | NM 198282.3 | ENST00000330794 | | 100 | | |
| TNFRSF13B | NM 012452.2 | ENST00000261652 | LRG 120 | 100 | | |
| TNFRSF13C | NM 052945.3 | ENST00000291232 | LRG 184 | 100 | | |
| TNFRSF1A | NM 001065.3 | ENST00000162749 | LRG 193 | 100 | | |
| TNFRSF4 | NM 003327.3 | ENST00000379236 | | 100 | | |
| TNFSF12 | NM 003809.2 | ENST00000293825 | | 100 | | |
| TPP2 | NM 003291 | ENST00000376065 | | 100 | | |
| TRAC | | ENST00000478163 | | 100 | | |
| TRAF3 | NM 145725.2 | ENST00000560371 | LRG 229 | 100 | | |
| TRAF3IP2 | | ENST00000340026 | | 100 | | |
| TREX1 | NM 033629.4 | ENST00000422277 | LRG 282 | 100 | | |
| TRNT1 | NM 001302946.1 | ENST00000251607 | | 100 | | |
| TTC37 | NM 014639.3 | ENST00000358746 | LRG 173 | 100 | | |
| TTC7A | NM 001288953.1 | ENST00000319190 | | 100 | | |
| TYK2 | NM 003331.4 | ENST00000525621 | LRG 121 | 100 | | |
| UNC119 | NM 005148.3 | ENST00000335765 | LRG 341 | 100 | | |
| UNC13D | NM 199242.2 | ENST00000207549 | LRG 122 | 100 | | |
| UNC93B1 | NM 030930.2 | ENST00000227471 | LRG 123 | 100 | | |
| UNG | NM 080911.2 | ENST00000242576 | LRG 124 | 100 | | |
| USB1 | NM 024598.3 | ENST00000219281 | LRG 352 | 100 | | |
| VPS13B | NM 017890.4 | ENST00000358544 | LRG 351 | 99.9992 | | |
| VPS45 | NP 009190.2 | ENST00000369130 | | 100 | | |
| WAS | NM 000377.2 | ENST00000376701 | LRG 125 | 100 | | |
| WIPF1 | NM 001077269.1 | ENST00000392547 | LRG 374 | 100 | | |
| XIAP | NM 001167.3 | ENST00000371199 | LRG 19 | 99.9986 | | |
| ZAP70 | NM 001079.3 | ENST00000264972 | LRG 126 | 100 | | |
| ZBTB24 | NM 014797.2 | ENST00000230122 | LRG 326 | 100 | | |
